# X-chromosome inactivation in human iPSCs provides insight into escape-regulated gene expression

**DOI:** 10.1101/2023.10.25.563960

**Authors:** Hande Topa, Clara Benoit-Pilven, Taru Tukiainen, Olli Pietiläinen

**Affiliations:** Neuroscience Center, Helsinki Institute of Life Science, University of Helsinki, Helsinki, Finland; Institute for Molecular Medicine Finland, Helsinki Institute of Life Science, University of Helsinki, Helsinki, Finland; The Stanley Center for Psychiatric Research at the Broad Institute, of MIT and Harvard, Cambridge, MA, USA

## Abstract

Epigenetic variation in the X chromosome inactivation (XCI) of human induced pluripotent stem cells (hiPSCs) can impact their ability to accurately model biological sex biases. However, the gene-wise landscape of escape from XCI remains unresolved in female hiPSCs. To characterize the patterns of escape from inactivation, we performed a systematic survey of allele specific expression (ASE) in 165 female hiPSC lines. The analysis revealed that the escape from XCI was non-random and affected primarily genes that escape also in human tissues. However, individual genes and cell lines varied in the frequency and degree of escape. The escape increased gradually after modest decrease of *XIST* in cultures, whose loss is commonly used to mark lines with eroded XCI. We identified three clusters of female lines at different stages of XCI. The increased degree of escape from XCI amplified female-biased expression and reduced male-female differences in genes with male-biased expression in the X chromosome. In autosomes, the increased escape modified genuine sex differences in a dose-dependent way suggesting that escape from XCI directly regulated autosomal gene expression. The variation in escape was sufficient to compensate for a dominant loss of function effect in several disease genes. The study presents a comprehensive view of escape from XCI in hiPSCs and emphasizes the need to monitor the XCI status at gene level for disease modeling. It further suggests that the uncommon and variable escape in hiPSCs can provide insight into X chromosome’s role in regulating gene expression and sex differences in humans.

## Introduction

There is great interest for using human induced pluripotent stem cells (hiPSCs) to understand cellular functions underlying male-female diversity in clinically relevant traits and disorders showing sex biases. Different lines can vary in their epigenetic landscape in the X chromosome, which can influence their utility especially for modeling cellular functions that lead to X-linked disorders^1–5^. However, little is known how X chromosome inactivation (XCI), a dosage compensation mechanism of gene expression, affects individual genes in female hiPSCs. XCI silences transcription from the extra copy of chromosome X in females (XX) to balance the expression dosage with that of XY males^6^. XCI in humans takes place during early embryonic development^7,8^. From there on, it is forwarded to all descending cells. XCI is random in humans, so that typically approximately half of the embryonic cells will pass on a transcriptionally active paternal copy and the rest will have an active maternal X chromosome^9^. Intriguingly, XCI does not equally cover the whole chromosome but more than 20% of genes are reported to be transcribed to varying degree also from the inactive copy of the X chromosome^10^. Escape from XCI can subsequently result in sex-biased gene expression and may contribute to phenotypic diversity in males and females^10,11^.

During reprogramming, hiPSCs undergo global remodeling of their epigenetic state, but they retain the same inactive and active copy of chromosome X as their somatic progenitor (Xi and Xa, respectively)^2,12^. Instead of being mosaic for Xi, however, hiPSC cultures are reported to be clonal suggesting that each line descends from a single somatic progenitor cell^1,2,12^. Nevertheless, hiPSCs display epigenetic variation including erosion of XCI in prolonged cultures^5^. The erosion is associated with the loss of expression of the long non-coding RNA *XIST*, which is required to achieve XCI^2,3,13^. Previously, low levels of *XIST* mRNA have been associated with loss of histone H3K27 trimethylation, demethylation of promoter CpGs and increased global bi-allelic expression from chromosome X consistent with XCI erosion^2,3,13^. However, less is known about the gene-wise patterns of XCI in hiPSCs and how individual genes are affected by loss of *XIST* expression. The variation in patterns of escape in hiPSCs can have important implications for biological modeling, for instance, by rescuing or inducing phenotypes associated with deleterious gene variants in X-linked developmental disorders^1^.

Here, we describe a systematic analysis of the landscape of XCI in hiPSCs and the gene-wise patterns of escape associated with silencing of *XIST*. We find that although hiPSCs had largely uniform patterns of XCI that were shared with human tissues, they displayed considerable line-to-line variation in the number and degree of escape of individual genes. The patterns of erosion of XCI, associated with the loss of *XIST*, were found to be non-random and resulted primarily from escape of specific subset of epigenetically variable genes, including genes that have a degree of escape also in human tissues. The increased escape in subset of lines amplified female-biased expression and decreased sex differences for male-biased genes in the X chromosome and modified genuine sex differences in autosomal gene expression. Importantly, several genes underlying developmental disorders escaped silencing in the hiPSCs, which could modify biological effects associated with gene variants if not properly accounted for in the study design. Our results highlight previously unappreciated gene-wise patterns of XCI in hiPSCs. The gene-wise variability in escape from XCI needs to be carefully monitored in hiPSC study design, but it can provide novel insights into genetic mechanisms regulated by escape from XCI in humans.

## Results

### hiPSC cultures were clonal and yielded unbiased ASE estimates from bulk RNAseq

To investigate gene-wise landscape of XCI in hiPSCs, we analyzed allele specific expression (ASE) from RNA sequencing data (RNAseq) in 282 karyotypically normal cell lines from the HipSci project (165 female, **Table S1**)^14^. Unlike human tissues, hiPSC lines are thought to be clonal for Xi enabling assessment of XCI through ASE analysis from bulk RNAseq data^1,2,12^. We computed ASE by calculating the total fraction of RNAseq reads that mapped to the allele with fewer sequence reads for each gene. In total, ASE was computed for 12,911 genes that had at least 20 reads overlapping heterozygous sites in one or more cell lines (**Table S2, Table S3**). Out of these, 411 genes mapped to the X chromosome (**Table S3**). We then identified the female lines that had lost *XIST* expression, associated with transcriptional reactivation of the Xi^2,3,13^, and would therefore have eroded XCI. We defined a threshold for low *XIST* mRNA level based on the range of expression in female human tissues in the GTEx consortium^15^ (log_2_CPM (*XIST*) < 1.5, **Figure S1**). We identified 47 low *XIST* female lines and 118 lines with appreciable or high expression of *XIST* (**Figure 1A**). While there was no difference in the median ASE for the 12,500 autosomal genes between the female lines with high or low level of *XIST* and 117 male lines (median ASE = 0.44 (95% CI: 0.32-0.50), **Figure 1B**), the female hiPSCs with low *XIST* level, had significantly higher median and mean ASE at X chromosome genes than females with abundant *XIST* (p-value_median_ = 4.60e-03 and p-value_mean_ = 3.40e-11, respectively, one-sided Wilcoxon rank-sum test; median ASE (low-XIST lines) = 0.049 [95% CI: 0.008, 0.229]; median ASE (high-XIST lines) = 0.012 [95% CI: 0.007, 0.022]; mean ASE (low-XIST lines) = 0.179 [95% CI: 0.163, 0.223]; mean ASE (high-XIST lines) = 0.101 [95% CI: 0.094, 0.123]), **Figure 1C, Figure S2A-B, Table S1**), consistent with erosion of XCI, in line with previous work^3^. Furthermore, the results confirmed that the loss of *XIST* and erosion of XCI had little or no affect on ASE in autosomal genes, as expected.

**Figure 1.**
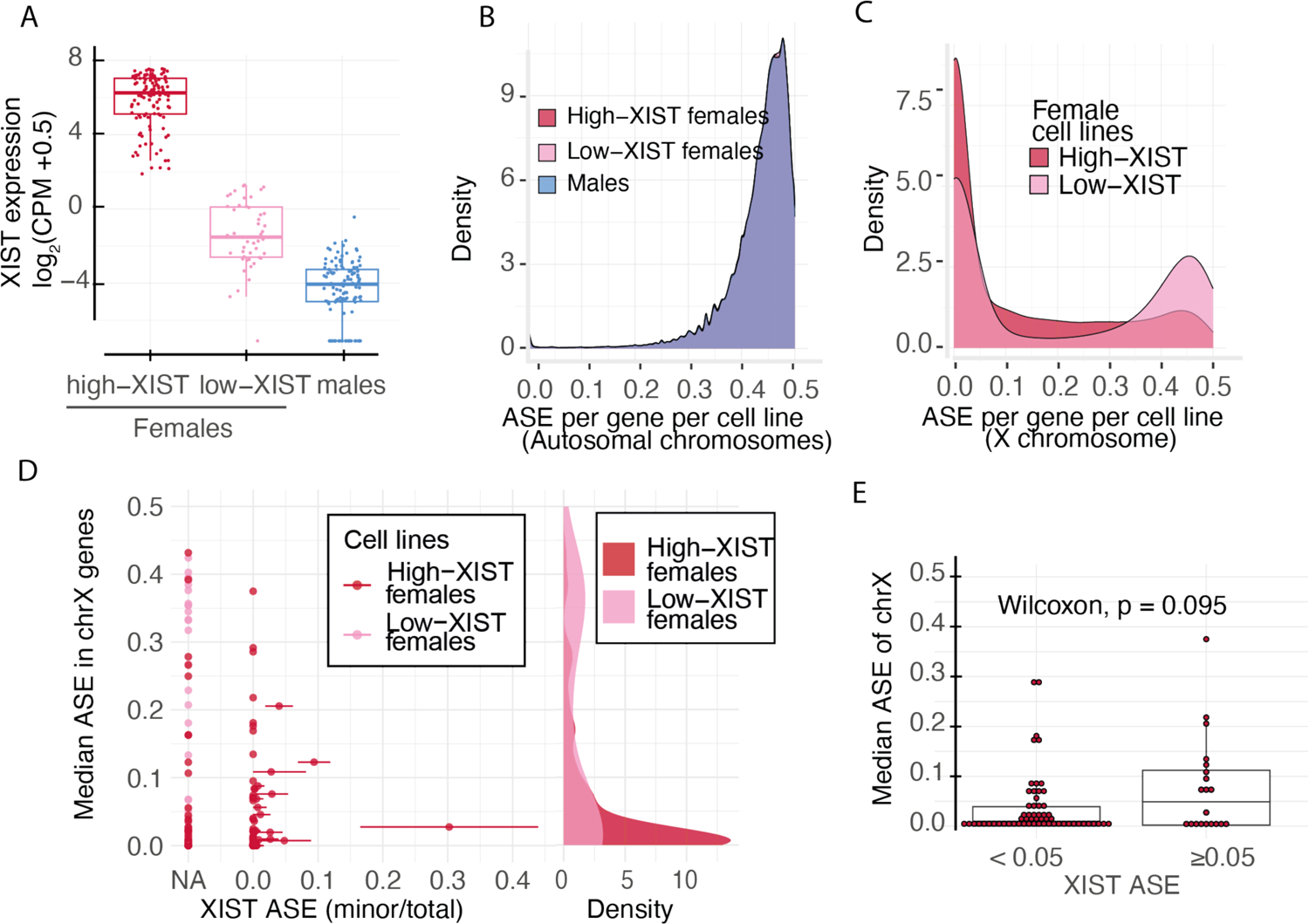
XIST expression is associated with escape from XCI. A) The hiPSCs were divided by sex and XIST expression to males and females with low or high XIST levels. B) The male and female lines with low or high level of XIST have similar ASE in autosomal genes. C) Female lines with low XIST levels differ in their median ASE from female lines with high XIST levels. D) The ASE for XIST could be defined for 80 lines. Majority of these lines (60) had monoallelic expression of XIST consistent with clonality of Xi in the cultures but varied in the level of escape from XCI, suggesting variable epigenetic patterns of the female hiPSCs. E) The 20 lines that had traces of biallelic expression of XIST (ASE ≥ 0.05), did not differ in median ASE of X-chromosome genes.

In addition to XCI erosion, loss of *XIST* expression is associated with naïve stem cell state with two active X chromosomes. Unlike the eroded lines, however, the naïve cells upregulate *XIST* upon differentiation to undergo XCI in culture^13^. To resolve between these two states, we compared *XIST* expression between 22 female hiPSCs and differentiated neuronal cells^16^ derived from the same hiPSC lines presumed to have the same passage number **(Table S1)**. We reasoned that an upregulation of *XIST* in the derived neurons compared to the hiPSCs could indicate a naïve state of the hiPSC from which the neurons were derived from. The analysis revealed only one female line with a notable upregulation of *XIST* upon differentiation, suggesting that most of the low *XIST* lines had undergone XCI and subsequently lost *XIST* during culture (**Figure S2C**).

Small number of female hiPSCs occasionally undergo reactivation of Xi during reprogramming that is followed by a subsequent random re-inactivation of one of the X chromosomes^5,17^. The resulting non-clonality can bias ASE estimates from bulk RNAseq data^18^. This is commonly observed for human embryonic stem cells^19,20^ but affects typically less than 10% of cells in hiPSCs^5,17^. To assess the impact of potential non-clonality for the ASE estimates, we analyzed allelic expression of *XIST*– transcribed solely from the Xi – to identify lines that were chimeric for Xi. We were able to obtain allelic information for *XIST* for 80 of the 165 female lines and found that most of the hiPSCs (75%) had monoallelic expression of *XIST* (ASE < 0.05), consistent with a clonal culture **(Table S1)**. In 20 lines, we observed a fraction of reads from the second allele of *XIST* that could reflect a re-inactivation of the other X chromosome. For all but one of these lines there was a significant allelic imbalance of *XIST* with over 90% of the reads mapping to only one of the alleles (**Figure 1D**), in line with the previous estimates^5,17^. However, the low level of bi-allelic expression of *XIST* was not significantly associated with increased median ASE (p-value = 0.09; Wilcoxon rank-sum test) or fraction of genes with ASE > 0.1 (p-value = 0.27; Wilcoxon rank-sum test) at the X chromosome (**Figure 1E, Figure S2D**). Together, these data are in line with previous estimates of clonality and single ancestor for each hiPSC line and suggest that the traces of non-clonality in the cultures do not significantly bias the ASE estimates from bulk RNAseq.

### Escape from XCI in hiPSCs affects distinct sets of genes

After confirming the clonal XiXa state of the hiPSCs and that we could accurately measure ASE, we sought to better understand the patterns of the XCI in the female lines. Beyond the differences in the mean ASE at X chromosome genes between the low and high *XIST* lines the ASE distributions were heavily biased towards monoallelic expression (ASE = 0) with long tails and bimodality (**Figure 1C**). This suggested that the degree of escape from XCI was varying by cell line and/or by gene in the female lines. To obtain, therefore, a clearer picture of the gene-wise patterns of XCI, we studied ASE of the 411 X chromosome genes that had allelic counts available in the 165 female lines. We found that 61% of the genes (n = 252) escaped in at least one cell line (ASE > 0.1; binomial test, FDR < 0.01, **Table S4**). This implied that a large fraction of genes had a potential to, on occasion, escape XCI in the hiPSCs. To understand whether the high rate of escape was explained by the loss of *XIST* in a subset of lines, we compared the number of genes with ASE > 0.1 (FDR < 0.01) between the female lines that had high or low levels of *XIST*. Surprisingly, we observed that a large number of genes escaped occasionally in both the high *XIST* and low *XIST* lines. Out of the 252 genes, 209 escaped in at least one line with high *XIST*, and 211 escaped in at least one line in the low *XIST* lines (168 escaped in both, **Table S5**). However, while the number of genes that escaped XCI in at least one line was similar in the two groups, the total fraction of genes that escaped XCI in each line was significantly higher in the female lines with low level of *XIST* than in lines with high *XIST* expression (p-value = 1.30e-08; t-test, high-XIST mean fraction: 0.27; 95% CI [0.244, 0.300], low-XIST mean fraction: 0.44; 95% CI [0.396, 0.488], **Figure 2A, Figure S3** panel vi**, Table S1)**. These results suggested that although lines with low level of *XIST* had on average more genes escaping per line, consistent with the erosion of XCI, many of the escaping genes had a propensity to sporadically escape XCI also in lines with high *XIST* levels.

**Figure 2.**
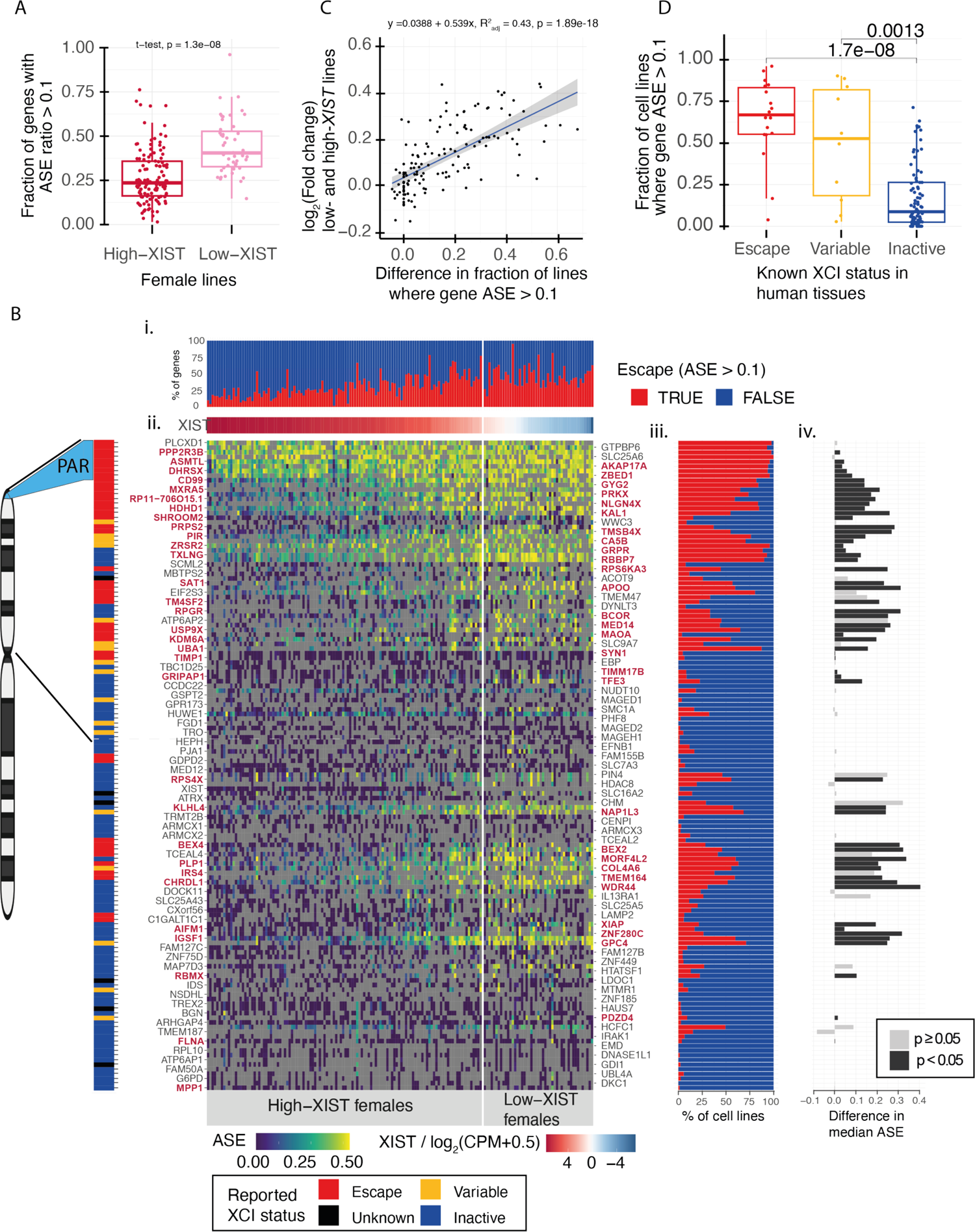
Landscape of escape from XCI in hiPSCs. A) A Tukey boxplot of fraction of genes with ASE > 0.1 in female lines with high and low level of XIST. Female lines with low XIST levels had significantly higher fraction of genes with ASE > 0.1, Wilcoxon p.value = 1.30e-08. B) Genomic pattern of escape from XCI for 139 X chromosome genes in 165 female lines. i) a bar graph of the percentage of escape (ASE > 0.1, FDR < 0.05) and inactive (ASE < 0.1) genes for each line. ii) A heatmap of degree of ASE for 139 genes (rows) in 165 female lines (columns). The genes are oriented according to position in X chromosome and the 165 lines are in decreasing order of XIST expression. The low XIST lines are separated with vertical line. Cells in grey color indicate missing information. The pseudo autosomal region (PAR) is highlighted in light blue in the chromosome. Names of the genes with significant difference in ASE between low XIST and high XIST lines (Wilcoxon rank-sum test, p < 0.05) are colored in red. The escape status in humans is color coded on the left bar for the genes for escape genes (red), variable (yellow), inactive (blue), black (unknown)). iii) The % of lines with inactive (blue) and escape (red) for ach gene is presented as a bargraph on the right side of the heatmap. iv) Difference in the degree of ASE between the low- and high-XIST female lines is presented as bar graph. The significant genes are highlighted in black bars. C) Difference in the fraction of lines where ASE > 0.1 between low XIST and high XIST female lines is positively correlated with the log fold changes between the gene expression levels of low XIST and high XIST female lines (p-value = 1.89e-18, r^2^ = 0.43). D) Boxplot of fraction of cell lines that have ASE > 0.1 of genes that are known to escape (red), variably escaping (yellow), and inactive (blue) in human tissues. Only the genes which have ASE data in at least 10 cell lines in both low-XIST and high-XIST female lines are included, excluding the PAR genes. The fraction of cells lines with ASE > 0.1 is significantly lower for inactive genes than for known escape and variably escaping genes (p = 1.70e-08 and 1.30e-03, respectively; one-sided Wilcoxon rank-sum test).

The large number of genes with occasional escape from XCI in hiPSCs prompted us to further investigate the genomic patterns of the escape along the X chromosome. In humans and mice, the escape from XCI is tightly regulated by early epigenetic confinement and spatial control of access to the DNA^10,21^. We were, therefore, interested in whether genes at specific loci were more likely to escape than others. To investigate the genomic patterns of escape from XCI, we focused on 139 X chromosome genes for which there were allelic counts from 10 or more lines from both the low and high *XIST* female groups (**Figure 2B, Figure S3**). This allowed us to analyze the likelihood for each gene to escape XCI across multiple female lines. Out of the 139 genes, majority (122) escaped XCI at least once in the 165 lines (ASE > 0.1, FDR < 0.01). However, both the fraction of genes that escaped in each line (1.8% – 95% of all genes with ASE data per line), as well as the frequency of lines that escaped at each gene were highly variable (0% – 100%, **Figure 2B**, panels i and iii). Notably however, most genes escaped in only few lines. Out of the 139 genes, 60 escaped in more than 25% of the lines and 39 genes escaped in over 50% of the lines. This indicated that although hiPSCs displayed considerable heterogeneity in the escape of individual genes, the likelihood for a gene to escape in hiPSCs was non-random, as some genes were more likely to escape than others.

The non-random pattern of escape was also clear in the female lines that had low *XIST* level. We identified 41 genes that had significantly higher fraction of cell lines with escape (p-value < 0.05, Fisher’s exact test) in the low *XIST* female lines than in the high *XIST* lines (**Table S5**, **Figure S3**, panel vi), suggesting that escape due to the loss of *XIST* was concentrated to a subset of genes. Escape from XCI typically leads to higher gene expression. Therefore, to confirm that the increased ASE in the female lines with low level of *XIST* resulted from excess escape from XCI, we analyzed differential expression of the 139 X chromosome genes (**Figure 4A**, **Figure S4, Table S6**). We found that the genes that escaped more frequently in low *XIST* female lines than in high *XIST* lines commonly also presented with significantly higher gene expression in the low *XIST* female lines (p-value = 1.89e-18, **Figure 2C, Figure S4**). Out of 41 genes that had significantly higher fraction of cell lines where the gene escaped (ASE > 0.1), 34 were also more highly expressed in low *XIST* lines than in high *XIST* lines (p-value = 2.21e-07; Fisher’s exact test). Taken together, these findings implied that some genes readily reactivate in the hiPSCs while others remain resistant to the erosion of XCI.

### Erosion of XCI affects predominantly genes that escape in humans

We next compared whether the genes that frequently escaped XCI in hiPSCs clustered to any specific chromosomal regions or escaped in human tissues^10^. Expectedly, we observed a high frequency of escaping at the pseudo autosomal region (PAR) in the tip of the p-arm of chromosome X in all lines, and there was no difference in frequency of escape between the high and low *XIST* females (**Figure 2B, Figure S3**). Beyond genes in the PAR, genes that are known to escape or that have variable escape in some human tissues escaped significantly more frequently in the hiPSCs than genes that do not escape in humans (p-value = 1.70e-08, p-value = 1.30e-03, respectively; one-sided Wilcoxon rank-sum test, **Figure 2D**). This implied that hiPSCs maintain patterns of somatic escape. In addition, we observed clusters of genes, such as *GRPR, TXLNG, RBBP7* and *DYNLT3, RPGR, and BCOR,* that are inactive in humans but frequently escaped in the hiPSCs. Recently deleterious mutations in *BCOR* were associated with reduced differentiation capacity in pluripotent stem cells^22^. Taken together, this suggested that some epigenetic marks underlying XCI may be subject to removal during reprogramming that have roles in the stem cell state and differentiation capacity of the hiPSCs.

Finally, we compared the patterns of escape from XCI between the female lines with high or low level of *XIST*. The increased escape associated with the loss of *XIST* clustered to regions of consecutive genes in the p- and q-arms of chromosome X (**Figure 2B**). In contrast, genes at other regions escaped in only a handful of lines. This implied that some regions were particularly sensitive to the loss of *XIST.* Remarkably, the genes in these regions were more likely to escape also in female hiPSCs with high *XIST* and these overlapped with known escape genes in humans. However, the female lines with low *XIST* had even higher frequency of escape for the known escape and variable genes, as well as for the inactive genes than the high *XIST* females (p-value = 1.50e-08 for escape and variable genes, p-value = 6.10e-13 for the inactive genes; paired Wilcoxon rank-sum test, **Figure S2E**).

### Loss of *XIST* is associated with higher degree of escape at genes that escape in humans

The escape from XCI rarely results in full expression from Xi in humans^10^. For this reason and given the observed large overlap of escaping genes between the females with high and low levels of *XIST*, we wondered whether the loss of *XIST* resulted in an increase in the degree of escape at individual genes. The analysis revealed 58 genes with significantly higher ASE (p-value < 0.05, Wilcoxon rank-sum test) in the females with low *XIST* level than in the high *XIST* group (**Figure 2B**, panel iv, **Table S5**). These genes overlapped with those that were found to frequently escape also in the high *XIST* females and with known escape genes. Importantly, the observed differences in ASE were not associated with technical differences between samples, such as the number of cell lines with allelic information **(Figure S3**, panel iv) or the average number of sequence-reads per gene (**Figure S3**, panel v). Taken together, these results demonstrated that the erosion of XCI due to loss of *XIST* is associated with increased degree of escape of genes that commonly escape also in human tissues and intact hiPSCs. This suggests that the mechanisms underlying the normative escape have a role in the escape associated with the erosion of XCI. Moreover, we noted an increase in the degree of escape with modestly decreased levels of *XIST* in the female lines, suggesting that the erosion takes place gradually in the hiPSCs.

### Erosion of XCI takes place in cultures with appreciable XIST expression

Low level of *XIST* mRNA is typically used as a marker for XCI erosion in female hiPSCs. The observed line-to-line variability in escape led us to further investigate the relationship between the *XIST* expression and the XCI erosion in the female lines. Fitting a LOWESS regression curve for the *XIST* level and the fraction of X-chromosome genes that had ASE > 0.1 revealed that the *XIST* expression had an inverse linear relationship with the escape, but only in lines where *XIST* mRNA was still abundantly present (**Figure 3A**). However, the relationship with *XIST* expression and fraction of genes escaping plateaued before the loss of *XIST*. This implied that although low *XIST* expression is informative of the XCI erosion in hiPSCs, the erosion takes place in the cells in the culture at stages where *XIST* expression is not yet exclusively indicative of the erosion.

**Figure 3.**
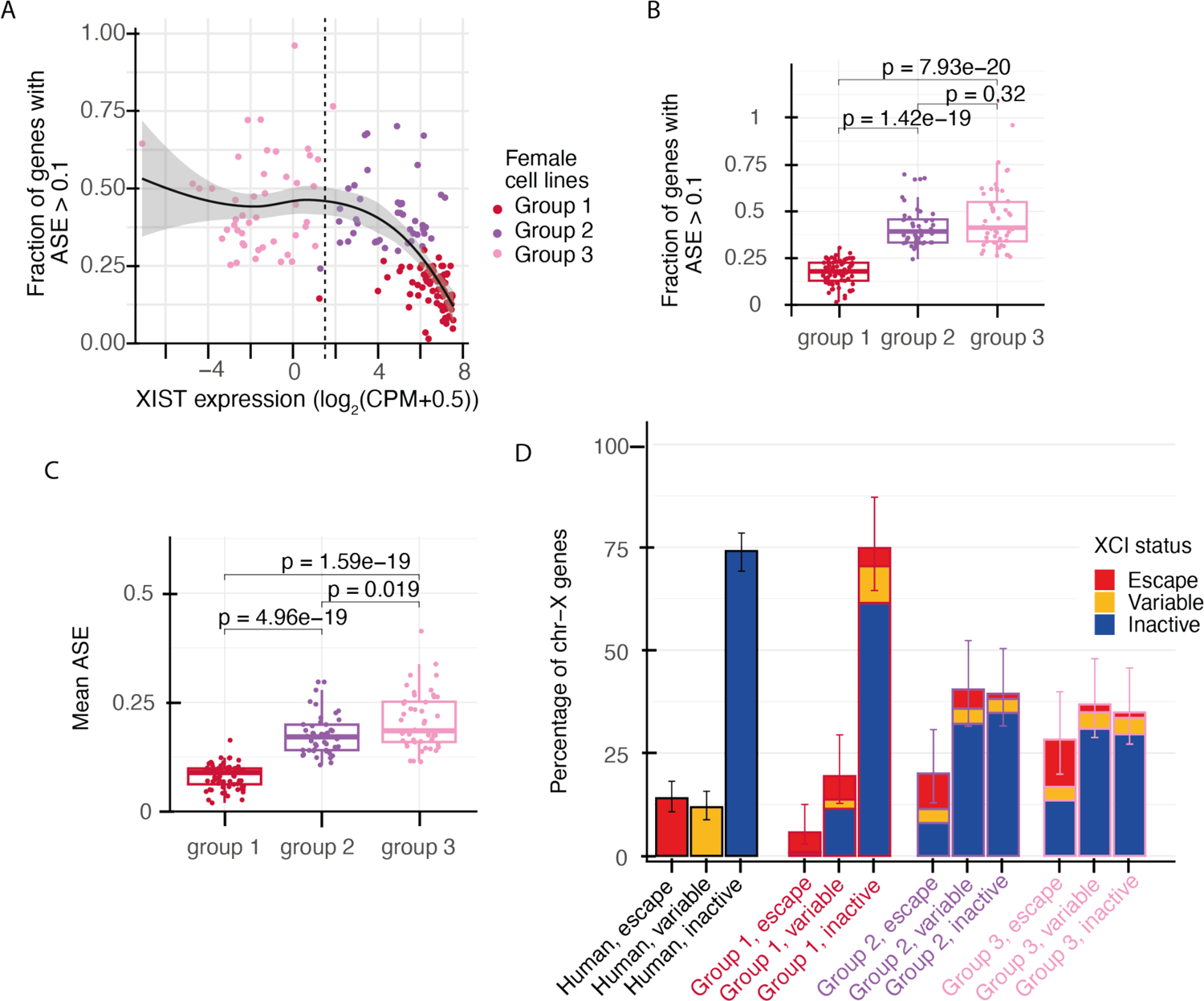
The XIST expression and fraction of escaping genes from XCI defines three groups of female lines. A) The relationship of XIST expression and fraction of X-chromosome genes that escape XCI (ASE > 0.1). K-means clustering of XIST level and fraction of genes with ASE > 0.1 identifies three groups of female lines. Group 1: high-XIST female lines with low level of escape (red), Group 2: Intermediate female lines that express XIST but have high fraction of escaping genes (violet). Group 3: Low-XIST females with high fraction of escape from XCI (pink). B) Fraction of genes with ASE > 0.1 is significantly different between group 1 females and group 2 and group 3 females, with the highest fraction of escaping genes in group 3 female lines (group 1 - group 2: p-value = 1.42e-19; group 2 - group 3: p-value = 0.32, group 1 - group 3: p-value = 7.93e-20). C) Mean ASE of X-chromosome genes is significantly different between the different groups of female lines with the highest mean ASE in group 3 female lines (group 1 – group 2: p-value = 4.96e-19; group 2 - group 3: p-value = 1.9e-02, group 1 – group 3: p-value = 1.59e-19). D) Group 1 female hiPSCs had similar percentage of inactive and variably escaping genes (at least one line with ASE > 0.1), and slightly less escape genes (>80% of lines with ASE > 0.1) than reported for human tissues (Chi-squared test for equality of proportions, p-value = 6.08e-05). The group 2 and group 3 female lines had significantly higher fraction of escaping and variable genes than in human tissues (Chi-squared test for equality of proportions, p-value = 5.75e-20 and p-value = 7.49e-22, respectively). In each of the female group comparisons with human tissues, only the genes which had available information in both of the groups are included.

The non-linear relationship of *XIST* expression and XCI erosion motivated us to define the state of XCI in female lines by combining information from *XIST* expression and the fraction of genes that escaped from XCI in each line. Therefore, we performed k-means clustering by using *XIST* mRNA levels and fractions of genes with ASE > 0.1. The optimal number of clusters was determined as three by the gap statistic method^23^ that revealed three distinct patterns of XCI (**Figure 3A**). Firstly, a group with high expression of *XIST* and low level of escape (0.18 median fraction of genes with ASE > 0.1) separated from the rest of the lines (high *XIST* group / group 1, n = 74). The remaining lines, clustered into two groups that had a significantly higher fraction of genes with ASE > 0.1 than the group 1 female lines. But while group 2 lines had an appreciable *XIST* expression (group 2 / intermediate group, 0.39 median fraction of genes with ASE < 0.1, n = 45), group 3 had a low level of *XIST* (0.41 median fraction of genes with ASE > 0.1, n=46), and corresponded to the original threshold for loss of *XIST* defined in the beginning of the study (**Figure 3B**). Although group 2 and group 3 had similar fraction of genes with ASE > 0.1 (p-value = 0.32; Wilcoxon rank-sum test), the overall degree of escape was significantly higher in the group 3 than in the intermediate group 2 lines (p-value = 1.90e-02; Wilcoxon rank-sum test, **Figure 3C, Table S1**), suggesting progressive reactivation of Xi in the cultures. We compared the percentage of genes that were inactive, escaped in >80% of lines (escape genes), and that had variable escape (<80 % of lines with escape) in hiPSCs to the precentages observed in human female tissues^10^. The comparison revealed similar fractions of variable and inactive genes and slightly smaller fraction of escape genes in the group 1 hiPSCs and human tissues (Chi-squared test for equality of proportions, p-value = 6.08e-05, **Figure 3D, Figure S2F**). This suggested that the group 1 lines, presumably reflecting the most pristine group of the hiPSCs, had intact XCI with high similarity to humans. In comparison, the group 2 and group 3 lines had significantly higher fraction of variably escaping genes, likely resulting from progressive erosion of XCI (Chi-squared test for equality of proportions, p-value = 5.75e-20 and p-value = 7.49e-22, respectively).

### The escape from XCI induces the expression of genes with female biased expression

Female hiPSCs with low levels of *XIST* have been shown to differ in gene expression from the female lines with abundant expression of *XIST^3,5^*. How variability in the escape from XCI affects male-female differences has been less studied, however. To investigate the effect of the degree of escape from XCI to sex differences in gene expression in the X chromosome, we compared the three groups of female lines to 117 male hiPSCs (**Table S6**). The analysis revealed a progressive, general upregulation of X chromosome genes in the group 2 and 3 female lines that had increased level of escape from XCI (**Figure 4D**, horizontal bars). This was also evident from a growing number of increasingly inclusive sets of X chromosome genes that had higher expression in the three groups of female lines than in males. We identified 124 upregulated genes in the pristine group 1, 262 in the intermediate group 2, and 353 in the low *XIST* group 3 lines (**Figure 4D**). Close to all upregulated genes in the group 1 females, were found significantly upregulated also in the group 2 and group 3 females (116 out of 124 genes, 94%, FDR < 5%). Similarly, of the 262 significantly upregulated genes in group 2, 97% (256 genes) were significantly upregulated also in group 3 females. Notably, the mean expression of the significantly upregulated genes that overlapped between the three female groups increased in the lines with a higher degree of escape. This resulted in significantly greater differences between males and the group 2 and group 3 female lines than between males and the pristine group 1 females (**Figure 4B**). Conversely, genes that had significantly lower expression in the group 1 female lines than in male hiPSCs, mostly had attenuated sex bias in the group 2 and group 3 females (**Figure 4C**). Out of the 58 downregulated genes in group 1 females, 37 genes were significantly different from males only in group 1 female lines. These included PAR genes with a known male biased expression in human (**Figure 4D**). Furthermore, we found 11 genes that had significantly lower expression in group 1 females than in males but were upregulated in group 3 female lines. Two of these (*CLCN4* and *MID1*) were PAR genes and the rest spread the p-arm (*SCML1, DMD, PLP2, PRICKLE3, USP27X, SHROOM4*), and q-arm (*EFNB1, HMGB3, L1CAM*) of X, suggesting another mechanism for the flipping of the male biased expression for these genes in the group 3 lines. In summary, the findings demonstrated that increased escape from XCI results in upregulation of X chromosome genes. This resulted in magnified sex differences in expression of genes that had female biased expression in group 1 hiPSCs and decreased effects of genes with male biased expression.

**Figure 4.**
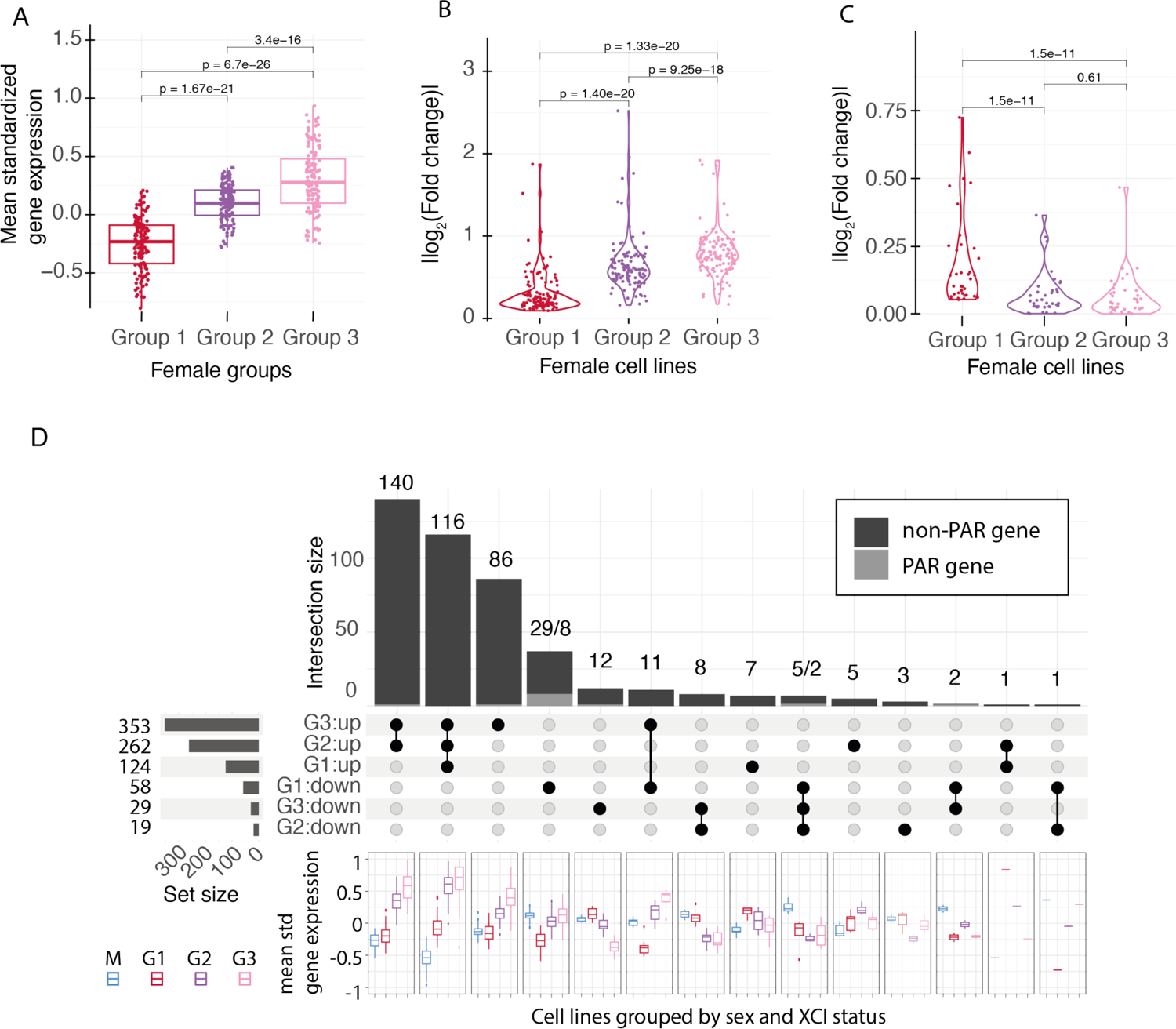
Escape from XCI enhances female-biased expression and reduces male-biased expression in X chromosome. A) Gene expression increased with higher escape from XCI for X chromosome genes (group 1 - group 2: p-value = 1.67e-21, group 2 - group 3: p-value = 3.40e-16, group 3 - group 1: p-value = 6.70e-26, paired t-test), a Tukey style boxplot. B) Violin plot of absolute log_2_(fold changes) for X chromosome genes with female biased expression in group 1. The fold changes increase significantly between group 1 and group 2 (p-value = 1.40e-20), group 2 and group 3 (p-value = 9.25e-18) and group 1-group 3 (p-value = 1.33e-20), paired t-test. C) A violin plot of X-chromosome genes with male-biased expression. The fold changes decrease significantly between group 1 and group 2 (p-value = 1.50e-11), group 1 and group 3 (p-value = 1.50e-11). There is no statistically significant differences between group 2 and group 3 (p-value = 0.61), paired t-test. D) An upset plot of the overlap of significantly differentially expressed X-chromosomal genes between male lines and the group 1 (G1), group 2 (G2) and group 3 (G3) female lines with different directions of effect. Boxplots below upset plot represent the mean standardized expression for males, and the group 1, group 2, and group 3 female lines for subsets of overlapping genes with significant male-female differences in expression.

### Escape from XCI is associated with dose-dependent sex differences in gene expression of autosomal genes

We next wondered whether the state of XCI in the female lines influenced male-female differences in the expression of autosomal genes. Differential gene expression analysis between the male and female lines revealed novel sex differences associated with the two groups with high degree of escape from XCI that were not present in lines with low escape. This was first evident from markedly higher number of significantly differentially expressed genes in lines with low *XIST* level and high degree of escape. We observed twice as many significant genes in the group 3 females (n=5,084 significant genes) than was detected in the low escape group 1 (n=2,364) or in the intermediate group 2 female lines (n=2,555, **Figure 5A**, pink, red, and violet horizontal bars, respectively). The disproportion in the number of significant genes was not driven by differences in power due to variable sample sizes; group 1 had the largest sample size (n=74) and the lowest number of differentially expressed genes (FDR < 5%), and groups 2 and 3 were equally sized (n=45 and 46, respectively). Instead, over half (54%) of the 5,084 significant genes were uniquely detected only in group 3 (n=2,729 genes, 1,428 downregulated and 1,301 upregulated, **Figure 5A**, pink vertical bars). This implied that prominent changes in gene expression took place in the lines with the highest degree of escape from XCI. Remarkably, the rest of the significant genes in group 3, were nearly completely shared by the intermediate group 2 females (86%, 2,018 out of 2,355 genes, **Figure 5A**, pink shaded, violet bars). These genes comprised the majority (79%) of the significant genes in group 2, demonstrating that beyond the excess of significant genes in group 3, the two groups with high fraction of escape genes shared marked similarities in sex biased gene expression.

**Figure 5.**
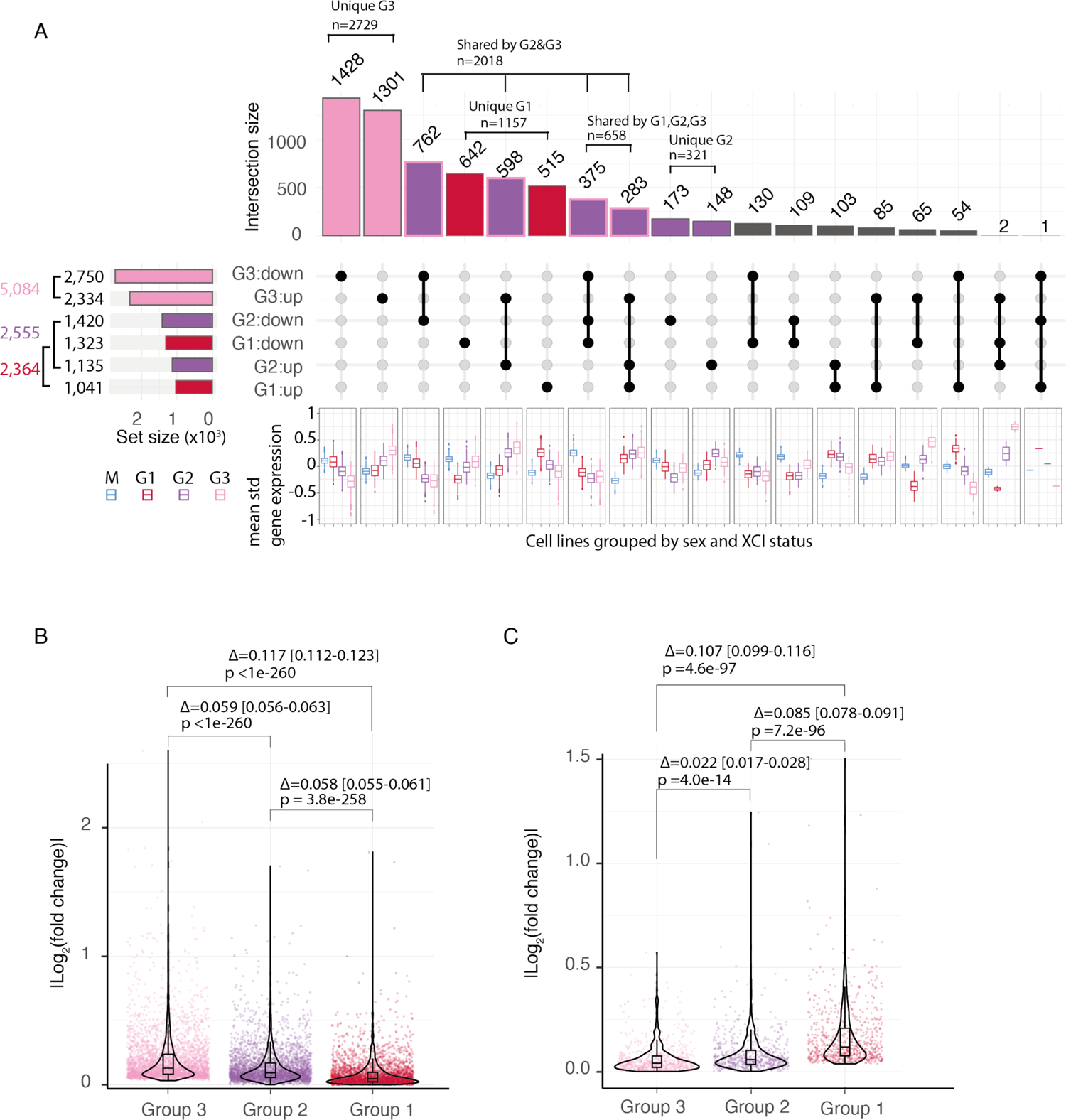
Differential expression of autosomal genes in male and female lines with different degree of escape. A) An upset plot for autosomal genes that were found significantly differentially expressed between males and the three groups of female lines. Boxplots below the upset plot present the mean standardized gene expression for male lines and the group 1 (G1), group 2 (G2) and group 3 (G3) female lines for subsets of overlapping genes similar to Figure 4D. The horizontal and vertical bars are colored by the female group: pink for group 3, violet for group 2 and red for group 1. The vertical bars indicate the sharing of significant male-female differences between the female groups. The unique effects in group 3 are colored in pink, the effects shared by group 2 and group 3 are colored in violet and outlined with pink, the unique effects in group 2 are colored in violet without outline and the effects that are unique for group 1 are colored in red. The non-concordant effects are colored in grey. B) A violin plot and embedded tukey style boxplot of the absolute log_2_ (fold-changes) for 3,260 genes significant in group 3 that have same direction of effect in all female groups. The fold changes associate with X chromosome dosage. The differences in absolute log_2_ (fold-changes) are shown above plots with associated p-values, paired t-test (p-value [group 3 - group 2] < 1e-260, p-value [group 3 - group 1] < 1e-260, p-value [group 2 - group 1] = 3.80e-258) C) A violin plot and embedded Tukey style boxplot of 672 genes significant in group 1 and same direction of effect in other groups. The differences in absolute log_2_ (fold-changes) are shown above plots with associated p-values, paired t-test (p-value [group 3 - group 2] = 4.00e-14, p-value [group 3 - group 1] = 4.60e-97, p-value [group 2 - group 1] = 7.20e-96).

In comparison, group 1 lines with high *XIST* and low escape had less overlap of significant genes with the other groups. Only 658 (33%) of the 2,018 common genes between group 2 and 3 were found significantly changed between males and the pristine group 1 females with intact XCI. This suggested that most of the male-female differences in the lines with high fraction of escape were weaker in group 1 or were aquired with advanced degree of escape, and, therefore, remained undetected in lines with low level of escape. In addition, there were 1,157 genes that were uniquely changed only in the group 1 females (49% of all significant genes in group 1, **Figure 5A**, red colored vertical bars). This implied that in addition to novel sex differences that emerged with higher degree of escape, some male-female differences that were observed in lines with low level of escape faded out due to increased escape from XCI.

The apparent high number of unique significant sex effects in group 3 and in group 1 lines prompted us to study more closely the patterns of mean expression of the different overlapping sets of significant genes between the groups of female lines (**Figure 5A**, boxplots). Intriguingly, we found that the mean expression of the genes with overlapping significant effects in the female lines followed a dose-dependent pattern with the degree of escape from XCI (**Figure 5A**). We noticed that for a large fraction of the overlapping sets of genes the mean expression in the intermediate group 2 lines was in between the levels of the low escape group 1 and the high escape group 3. Furthermore, there were only 173 downregulated and 148 upregulated genes that were unique for group 2 and whose mean expression was the lowest and the highest in group 2 with intermediate degree of escape, respectively (**Figure 5A**, violet colored vertical bars). This suggested that the X chromosome dosage was directly associated with the level of gene expression of genes in the autosomes and that they followed a dose-dependent relationship with the escape from XCI.

We then studied the concordance of the male-female differences of individual genes in lines with different degree of escape from XCI. The analysis revealed overlap of similar sex biases in the groups with high escape, whose magnitudes increased with the X chromosome dosage. In addition to the 2,018 shared significant genes in group 2 and group 3, 95% of the unique significant effects found only in group 3 were regulated to the same direction also in group 2 (n=2,581 out of 2,729), underlining overall similarity in the high escape groups. The concordance of the effects decreased slightly in the low escape lines in group 1 to 86% for the 2,018 shared genes by group 2 and group 3, but confirmed that the sex biases detected in the high escape lines had mostly similar effects in the low escape lines as well. Intriguingly, this sharing of effects lowered markedly for the 2,729 genes that were detected only in group 3 (56%), revealing only modest, but significant 1.1-fold change in the overlap beyond chance (p-value = 4.50e-05; binomial test). The smaller overlap suggested that effects were substantially weaker in group 1 or were acquired only in lines with more escape and previously inactive genes escaping XCI.

Next, we tested whether the decrease in the overlap of differentially expressed genes resulted from systematic quantitative differences in the magnitudes of the sex biases with different X chromosome dosage. For this, we compared the absolute fold changes associated with sex-biased expression in the subsets of genes with different overlapping effects across the female groups. We found that the magnitude of the male-female differences was associated with the degree of escape from XCI in the female hiPSCs. We combined genes with concordant same-direction of effects in the three female groups and compared the magnitudes of associated absolute fold-changes. The analysis revealed the largest absolute difference from males in the group 3 lines with the highest degree of escape. The magnitudes decreased significantly with reduced X chromosome dosage in the intermediated group 2 and the lowest average effects were observed in group 1 (**Figure 5B**). Importantly, the dose-dependent trend was detectable in most genes. Out of the 3,260 genes with concordant effects, 80% had the largest absolute fold change in group 3 and 83% the lowest difference from males in group 1, confirming that the averages were not driven by a small subset of genes. Moreover, the quantitative dose-dependent trend was observed for each subset of overlapping significant genes detected in group 3 and group 2 (**Figure S5A-C**). In addition, we found that the 2,729 genes, significant only in lines with the highest degree of escape, had on average smaller effects than the genes that passed the threshold for significance already in the intermediate group 2 and group 1 (**Figure S5D**). This further confirmed that the effect sizes of the increasingly inclusive subsets of genes detected in the female lines with increasing degree of escape were associated with the X chromosome dosage. Together these results implied that the progressive gene expression differences were quantitatively associated to the X chromosome dosage in the female hiPSCs, pointing to a potential direct mechanism to X chromome regulated autosomal expression.

Finally, we compared the effects of 1,157 genes that were uniquely detected only in the low escape group 1 lines. Out of these, 672 had same direction of effect in all groups. Conversely to the increasing magnitudes of female biased expression associated with the genes detected in group 2 and group 3, the genes unique to group 1 followed an opposite dose-dependent trend with increased escape from XCI. For these genes we found the lowest overlap of same direction of effecs in group 3 (61%) and with lowest average absolute fold changes (**Figure 5B**). These results demonstrated that in addition to enhancing sex biases in gene expression, the increased escape resulted in fading out of some male-female biases in gene expression. This was similar to what was observed e.g. for genes in the PAR of X chromosome that in high escape female lines became more similarly expressed with males than in females with intact XCI, and could imply a role for these genes regulating male-biased expression in the autosomes.

### Genes underlying X-linked developmental disorders vary in escape in hiPSCs

Previously, erosion of XCI has been reported to mask genetic effects related to X-linked developmental disorders^1^. This led us to systematically explore how line-to-line variability in escape from XCI impacts the expression of genes where gene disabling variants have been previously implicated in neurodevelopmental disorders (NDD) both in X chromosome and autosomes. We gathered 458 genes previously associated in neurodevelopmental diseases (NDD)^24–30^. We identified, 78, 114, and 184 NDD genes that were significantly (FDR < 5%) differentially expressed in the group 1, group 2, and group 3 females compared to males, respectively. (**Table S6**). A likely primary genetic mechanism for a dominant loss of function (LoF) variant is through the loss of a functional copy of a gene and reduced gene expression^31^. We, therefore, focused on genes whose 95% confidence interval of absolute fold-change (FC) covers 1.5, corresponding to 50% increase or decrease in gene expression. We reasoned that a degree of escape comparable to gaining an extra gene copy would be sufficient to mask the predicted disease-mechanism of reduced expression associated with LoF variants^12^. We identified 32 NDD genes that had an absolute FC of ≥ 1.5 and were significantly differentially expressed in at least one of the female groups (Group 1, Group 2, and Group 3) compared to males. Out of these, 17 were X-linked and 15 were autosomal (**Figure S6**). These results indicated that the escape from XCI may influence also the penetrance of autosomal variants. ASE information could be obtained for 6 of the chromosome X genes in the female lines, each displaying higher ASE ratio in the low-XIST lines (**Figure 6A,B**), further confirming that the increased expression resulted from escape from XCI for these genes as a direct compensatory mechanism. These genes demonstrated a dose-dependent increase in expression associated with the higher degree of escape. Importantly, however, some of the genes, like *KDM6A* and *TFE3*, displayed considerable line-to-line variation in expression and ASE. This suggested that using *XIST* expression alone may not be sufficient for accounting the variability in XCI in disease modeling.

**Figure 6.**
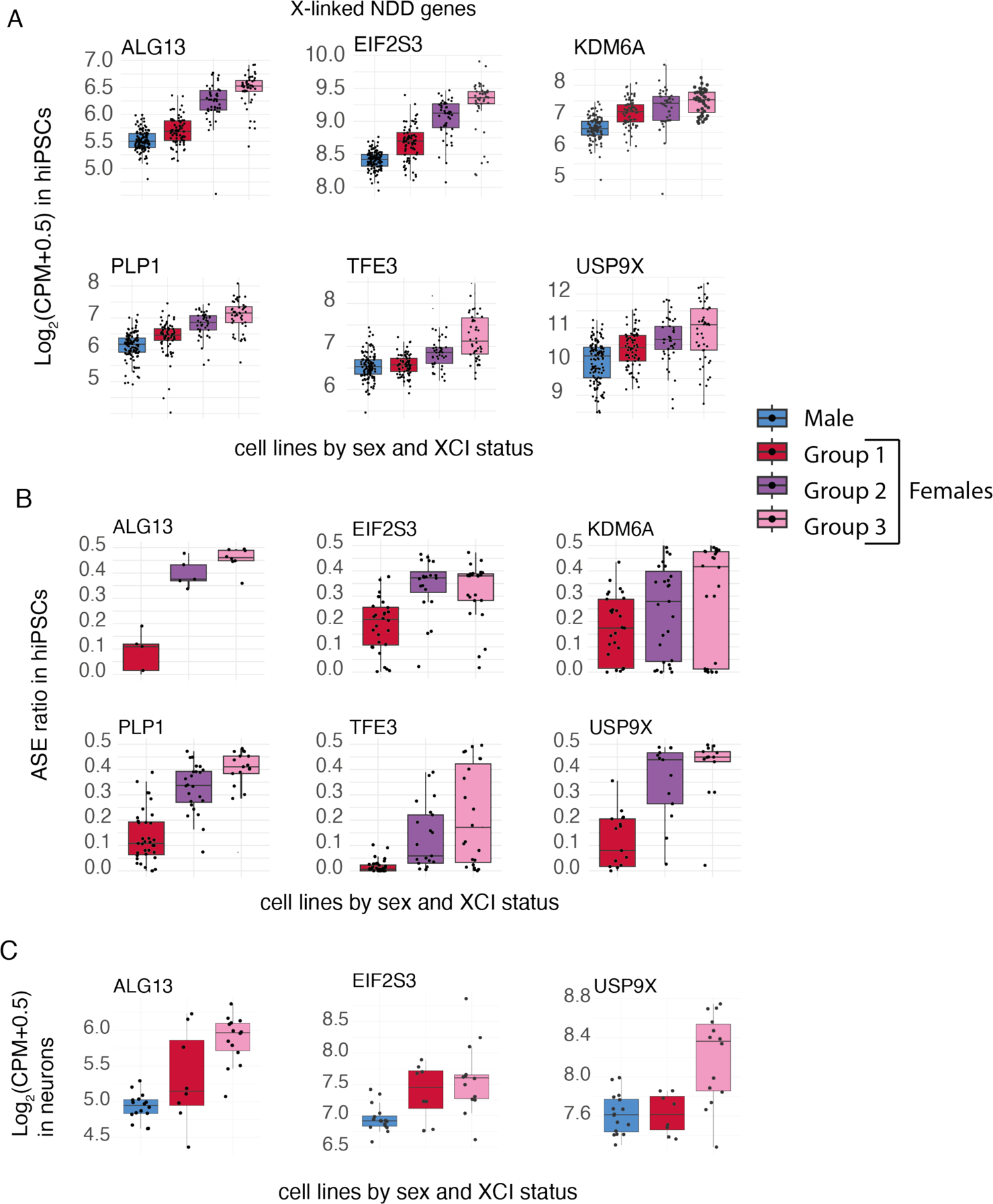
Variable escape from XCI in the hiPSCs masks dominant loss of function effects of genes in X-linked neurodevelopmental disorders (NDD). A) Boxplots of expression of selected genes linked with X-linked NDD with ASE and differential expression data that have significantly induced expression in group 3 female lines. B) Boxplots of median ASE for the disease genes show higher ASE in group 3 female lines. C) Boxplots of gene expression in differentiated neurons with high or low expression of XIST differentiated from the same hiPSC. Boxplots are in Tukey style.

Given that cell type specific pathogenic mechanisms associated with NDD genes are not necessarily present in hiPSCs, we asked whether escape from XCI in the differentiated neurons from the same hiPSCs resulted in compensatory gene expression of these genes. The analysis in the differentiated neurons (16 male, 8 high-XIST female and 14 low-XIST female) revealed 16 NDD genes (10 in chromosome X and 6 autosomal) which can reach 50% increase in gene expression in low-*XIST* female neurons (**Figure S7, Table S7**). Out of the six genes with ASE data available in hiPSCs, three were present in neurons and displayed similar patterns of expression with loss of XIST as observed in hiPSCs (**Figure 6C**). These results implicated that variation in XCI can directly reduce penetrance of gene disabling variants in hiPSCs and in differentiated cell types, including neurons, and the degree of escape of individual genes needs to be considered in biological modeling of X-linked diseases.

## Discussion

Understanding the gene-wise patterns of escape from XCI and its influence on sex differences are important for implementing correct study designs for hiPSC-based biological modeling. However, the landscape of XCI has not been systematically surveyed in large number of lines. Here, we performed a systematic analysis of gene-wise patterns of escape from XCI in 165 female hiPSCs using the HipSci resource^14^. The careful analysis of ASE across number of female lines expands the previous knowledge of erosion of XCI and provides novel insight into genomic patterns of escape in intact and eroded hiPSCs. Furthermore, by identifying groups of lines with different number and degree of escape from XCI, the data yields mechanistic insight into escape from XCI and its impact in regulating genetic sex differences in human cells. With the growing interest for mechanisms of sex differences in the field^32^, these data highlight the possibility to use the uncommon escape patterns of hiPSCs to understand how escape from XCI regulates the rest of the genome.

Several previous works have demonstrated global epigenetic variation in hiPSCs that is associated with the irreversible loss of *XIST* expression that leads to the erosion of XCI in some lines^2,3^. These studies have suggested screening cell lines for markers of XCI, including the loss of *XIST* expression to benchmark the quality of the cells. Here, the analysis of escape at the level of individual genes provides a more nuanced view to the erosion of XCI. We found that the fraction of genes with a degree of escape increased in lines prior to a considerable decrease in the *XIST* expression in the culture. These high-XIST high escape lines had similar fraction of genes escaping XCI as the lines with lowest XIST level. However, with the loss of XIST the degree of escape for these genes increased significantly. These findings are consistent with cell-level heterogeneity in the lines and suggested that XCI erosion progresses gradually to affect a growing number of cells in culture. These results underline the importance of applying genome-wide analysis for accurate and sensitive classification of XCI state in the hiPSCs, as suggested previously^4^.

In humans, XCI is tightly regulated, and different tissues generally have uniform patterns of escape from XCI^10,21^. We found that intact hiPSCs recapitulated the somatic landscape of escape^10^. The data further revealed that the escape associated with erosion of XCI was, similarly, non-random and affected regionally clustered genes along the X chromosome. These included a significantly higher degree of escape at genes that escaped in intact hiPSCs and in humans^10^. Previously, studies in mouse suggested that the reactivation of XCI during iPSC reprogramming takes place in a hierarchical order so that genes that escape XCI early are located near genes that escape XCI^33^. The escape from XCI is controlled by chromatin accessibility in the Xi. *XIST* expression is required for the formation of two densely packed super domains in the Xi with only regions of escaping genes preserving local structure of topologically associated domains with accessibility to the DNA^34^. Our data suggest that similar regional mechanisms that make some genes susceptible for the normative escape from the XCI in human tissues may regulate the erosion of XCI as well. However, more studies are needed to understand the exact mechanism that make some genes more likely to escape XCI than others.

One of the key insights revealed by the study is how variability in escape from XCI affected male-female differences in gene expression. The dose-dependent association between the degree of escape and the magnitude in the associated male-female differences in gene expression at autosomes implied that escape from XCI directly regulated the expression of these genes. This underlines, that the uncommon and variable escape in hiPSCs – often considered disadvantageous – could yield mechanistic insights into sex effects. Nevertheless, the variability in escape impacted the expression of numerous genes both in the X chromosome and autosomes. The variability in escape from XCI has been previously associated with key properties of hiPSCs, such as the differentiation ability^5,35^. Intriguingly, we found that *BCOR,* an inactive gene in humans where deleterious mutations are associated with increased proliferation and reduced differentiation ability^22^, escaped readily in female hiPSCs suggesting that the escape could be advantageous for the stem cell properties of the cells. Conversely, we found several dominant disease-linked genes whose expression was changed by escape from XCI and suggest that these modifying effects can complicate biological modeling if not addressed appropriately. Exciting progress is made to prevent and restore *XIST* expression and for the de-erosion of hiPSCs^36,37^. However, until such methods are readily available, it remains critical to account for the variability through careful experimental design.

## Limitations

The extreme skew of XCI resulting from the clonality of Xi in the hiPSCs enable detailed ASE analysis of X chromosomal genes using high-coverage bulk-RNA sequencing. However, it is important to note that without single cell resolution we cannot address the variability between individual cells in a culture. Therefore, the observed differences in ASE and gene expression may result from quantitative changes that take place in all cells in a culture, or larger changes that occur only in a subset of all cells in a culture. In addition, although we show extreme variability between hiPSC lines in fraction of genes with escape, we cannot completely rule out a possibility that some ASE estimates are affected by traces of non-clonality in the cultures. We used allelic expression of *XIST*, only expressed from the same Xi in clonal cultures, to verify clonality of the cultures. We observed a handful of lines with small degree of bi-allelic expression of *XIST*. However, this was not significantly associated with the median ASE and is therefore unlikely to significantly bias the results. Furthermore, the extreme allelic imbalance in *XIST* for all lines is consistent with a single somatic cell origin for the lines. With the current data set, we are unable to distinguish whether the bi-allelic expression of *XIST* results from non-clonality of the line, sequencing error, or other epigenetic variability in the cultures, such as transcriptional activation of *XIST* in the Xa. Importantly, ASE ratio of *XIST* could only be measured for a subset of cells with sufficient *XIST* mRNA levels and heterozygous coding variants in *XIST*. Therefore, we are unable to confirm the clonality of all lines, and the interpretation of the results is grounded on an assumption – strongly supported by the data and previous reports – that lines with low and high-levels of XIST are similar in terms of clonality of the cultures.

## Acknowledgements

We warmly thank the HipSCi project for rich iPSC data resource utilized in this study. The work was supported by funding for O.P. from the Instrumnetarium Science Foundation, Jalmari and Rauha Ahokas Foundation, Päivikki and Sakari Sohlberg Foundation, Jenny and Antti Wihuri Foundation, and Niilo Helander Foundation. T.T. was funded by the Research Council of Finland (grant numbers 315589 and 345867), Sigrid Jusélius Foundation, University of Helsinki Funds, and the HiLIFE Fellows Program.

## Author Contributions

The project was conceived and planned by H.T., C.B.P., T.T. and O.P. The analysis was performed by H.T. and assisted by C.B.P. with supervision from O.P. and T.T., O.P. and H.T. wrote the manuscript and it was commented and edited by C.B.P. and T.T.

## Supplementary tables

**Table S1.** Meta-data for HipSci hiPSC lines and differentiated sensory neurons used in the study as well as per-line ASE summary data of chromosome X genes for female hiPSC lines.

**Table S2.** ASE matrix for autosomal genes in iPSC lines.

**Table S3.** ASE matrix for chromosome X genes in female iPSC lines.

**Table S4.** Boolean matrix of escape (True / False) for X chromosome genes in female hiPSCs.

**Table S5.** Per-gene ASE summary data of female hiPSC lines for chromosome X genes.

**Table S6.** Differential expression analysis results for hiPSCs including comparisons for Males vs. Group 1 females, Males vs. Group 2 females, Males vs. Group 3 females and high-XIST females vs. low-XIST females.

**Table S7.** Differential expression analysis results for differentiated sensory neurons including comparisons for Males vs. high-XIST females and Males vs. low-XIST females.

## Methods

### Data Set

The data used in this study has been previously generated by the HipSci project^14^ and is available at https://www.hipsci.org/. This study included 282 cell lines (117 male and 165 female) that were derived from the skin tissues of healthy donors with predicted population of European, grown in feeder-free culture, and had the open access data available at the HipSci portal for RNA-seq and Exome-seq analyses (**Table S1**). The RNAseq fastq files, variant call files (vcf, whole exome sequencing mpileup variant calls), and RNAseq bam files (RNAseq mapped using splice-aware STAR) were used in ASE analysis, and can be accessed with study accession numbers PRJEB7388 (RNAseq) and PRJEB7243 (the whole-exome sequencing mpileup variant calls) in European Nucleotide Archive (ENA). Only fastq files from one sequencing center per cell line were include in the study. The fastq files for sensory neurons differentiated previously from 38 (16 male and 22 female) of the hiPSC lines^16^ were downloaded from ENA (PRJEB18630). Information about the differentiated sensory neurons used in the study are also listed in **Table S1**.

### RNA-seq data processing

RNA-seq data processing pipeline which is similar to the one used by GTEx project was used (https://github.com/broadinstitute/gtex-pipeline/blob/master/TOPMed_RNAseq_pipeline.md). More specifically, Homo_sapiens_assembly38.fasta (GRCh38) is used as reference genome, and ALT, HLA, Decoy contigs were excluded from the reference genome fasta file using the Python code provided in the GTEx pipline. The comprehensive gene annotation file gencode.v26.annotation.gtf was downloaded from https://www.gencodegenes.org/human/release_26.html, and all isoforms of a gene were combined into a single transcript using the collapsed gene model as explained in the GTEx pipeline at https://github.com/broadinstitute/gtex-pipeline/tree/master/gene_model (collapse_annotation.py). Prior to this, salmon package^38^ was used to confirm that HipSci lines had been sequenced using an unstranded protocol, similar to the GTEx samples.

The Illumina adapter sequences were trimmed using trimmomatic version 0.39^39^ and the trimmed fasta files were aligned to the reference genome using STAR version 2.6.0c^40^. Gene counts were estimated using RNA-SeQC version 2.3.5^41^. The gene counts from technical replicates were summed together for each cell line. The same RNA-seq data processing pipeline was carried out for the hiPSC lines and the differentiated sensory neurons.

### Differential gene expression analysis

Differential expression (DE) analysis was conducted on normalized gene counts for protein-coding genes and long intergenic noncoding RNAs (16,831 and 15,369 genes in total for hiPSCS and differentiated sensory neurons, respectively). For hiPSCs, we performed two separate DE analyses: one in which females were divided into two groups (low *XIST* and high *XIST*) and one in which females were divided into three groups according to *XIST* expression and fraction of escape genes (Groups 1, 2, 3). For differentiated sensory neurons, the DE analysis was performed for females with low *XIST* and high *XIST*. In each analysis, the genes that had less than 10 counts were filtered out. A TMM normalization was used for the gene-wise read counts and converted to log_2_(CPM+0.5) in the limma-voom package^42^. In addition, surrogate variables, which were adjusted for in the DE analyses, were estimated by the sva R package^43^.

For the DE analysis a mixed linear frame-work, implemented in the DREAM method^44^ was used that allowed accounting for the correlation between multiple replicate samples obtained from the same donor. The DE analysis for the sensory neurons included only one replicate sample per each donor and limma-voom^42^ was used for the DE analysis with the edgeR R package^45–47^. The DE analyses for hiPSCs included the following comparisons: high-XIST females vs. low-XIST females, males vs. Group 1 females, males vs. Group 2 females, males vs. Group 3 females (**Table S6**), and for differentiated sensory neurons: males vs. high-XIST females, males vs. low-XIST females (**Table S7**). Genes with adjusted p-value < 0.05 were considered as differentially expressed in each comparison. All statistical analyses were performed using R Statistical Software (v4.0.5)^49^. Figures were produced by using ggplot2^50^ and ComplexUpset^51,52^ libraries.

### Allele specific expression (ASE) analyses

To estimate the read counts which mapped to biallelic variants in each hiPSC line, the Genome Analysis Toolkit GATK^48^ was used with the HipSci vcf (exome-seq mpileup) and RNAseq bam files. For each line, GATK (version 4.2.0.0) ASEReadCounter was run with options-min-mapping-quality 10 and –min-base-quality 2, using the hs37d5 assembly and GencodeV19 annotation of the human reference genome. All biallelic variants that had at least 20 reads mapped to them were included to further ASE analysis. When calculating gene-level ASE, among all the variants located within a gene, only the variants which possessed the bigger count size if there was a gap less than the read length (100 bp) between any two variants were considered. The alleles which had the smaller number of counts (minor alleles), with respect to their counterparts were assumed to be located on the same chromosome haplotype. ASE was calculated for each gene by dividing the total number of reads that mapped to the minor alleles with the total number of reads that mapped to all alleles. Only the biallelic variants located along the gene were indlcuded in the analysis. ASE was calculated for 12,500 autosomal and 411 X chromosome genes in at least one of the hiPSC lines. For each X chromosome gene per line, alternative hypothesis of ASE > 0.1 was tested using binomial test to determine whether the gene escaped XCI (FDR < 0.01). The clonality of cell lines was tested by introducing a stricter ASE threshold (ASE < 0.05) for the *XIST* gene. *XIST* was defined to be monoallelicly expressed if the null hypothesis of ASE ≥ 0.05 was rejected for the *XIST* gene. It is important to note that the RNA-seq data was not realigned with a SNP-aware method, which may cause a bias toward the reference allele in the ASE results.

### Grouping of female lines by escape from XCI

A threshold for normative *XIST* expression was defined from *XIST* expression in human tissues from GTEx_Analysis_v8 results (available at https://gtexportal.org/). A threshold for loss of *XIST* was defined as having lower *XIST* mRNA level than observed for anyt tissues in GTEx (< 1.5 log_2_CPM). The *XIST* expression threshold of 1.5 log_2_CPM was used to divide the female hiPSCs as well as the female differentiated sensory neurons into two groups as low *XIST* and high *XIST* females. By combining ASE information with *XIST* expression level, a more refined grouping of female hiPSCs was obtained. For this, a k-means clustering was performed using the standardized fraction of genes with ASE > 0.1 and the standardized *XIST* expression levels. The gap statistic method^23^ suggested an optimal number of three clusters named Group 1, Group 2, and Group 3 females.

---

**Figure S1.**
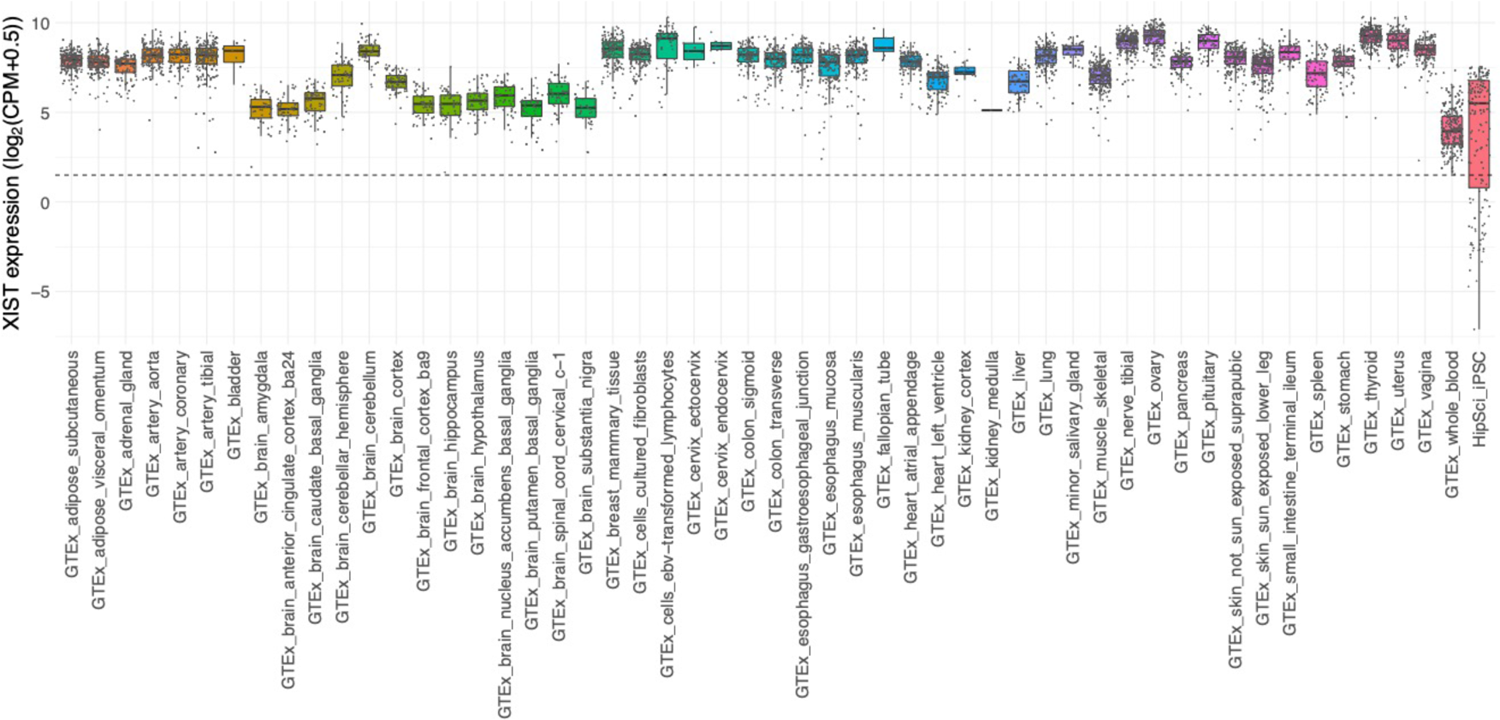
XIST expression in female human tissues in GTEx and hiPSCs. All female GTEx tissues had an XIST expression level of at least 1.5 log_2_CPM (indicated by the dashed horizontal line). Female hiPSCs displayed a wider range of XIST expression ranging below 1.5 log_2_CPM that was used to define a threshold for low XIST.

**Figure S2.**
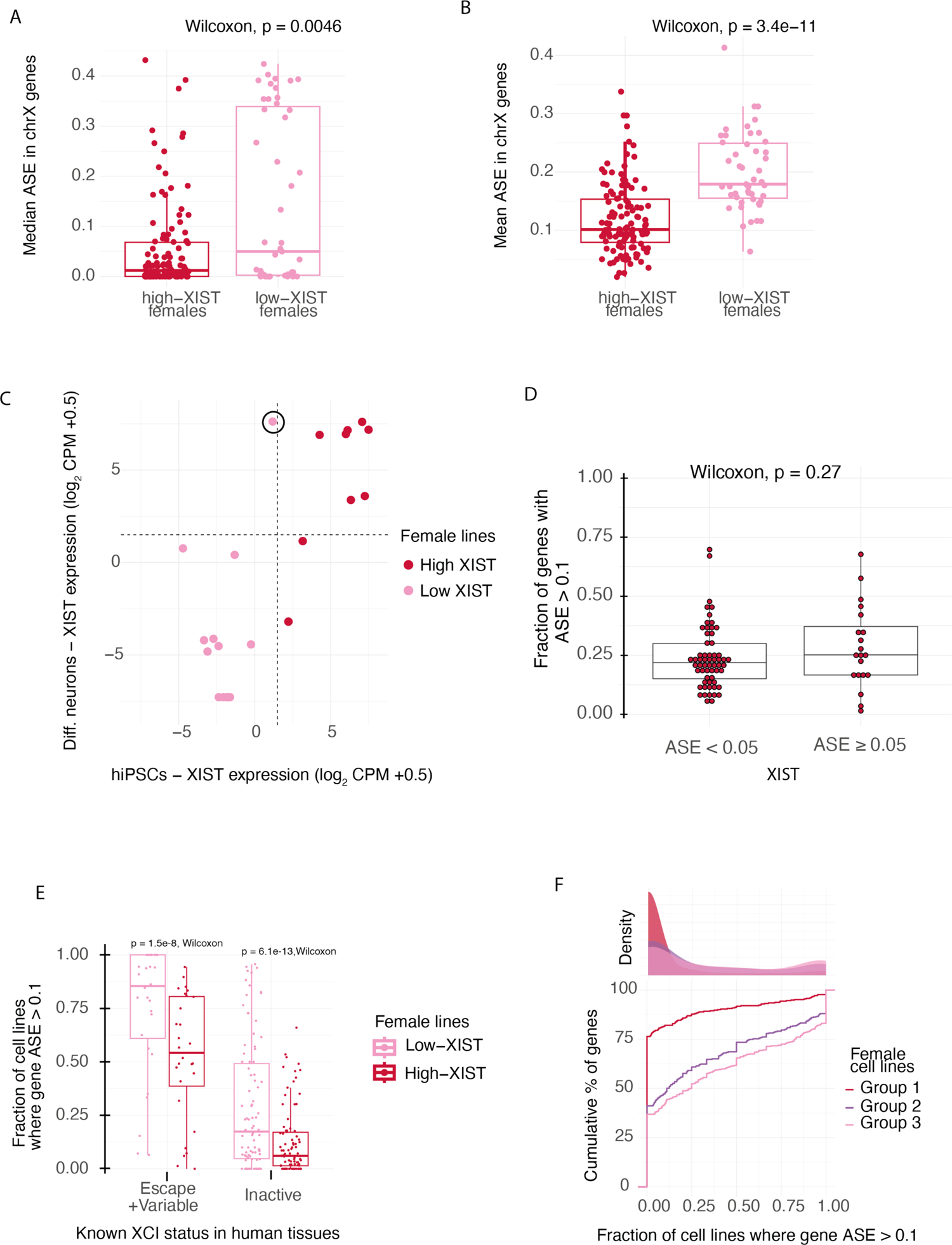
ASE of X chromosome genes in high and low XIST female lines. A) Median and B) mean ASE of X chromosome genes in female hiPSC lines with high or low level of XIST (n=118 and 47, respectively). C) XIST expression in hiPSCs and differentiated sensory neurons show high similarity with exception of one line (circled). D) Fraction of genes with ASE > 0.1 does not differ between lines with monoallelic expression of XIST (ASE < 0.05) or traces of bi-allelic expression (ASE ≥ 0.05) of XIST. E) A boxplot of fraction of cell lines where gene ASE > 0.1 in genes that are known to escape in humans and that are inactive in humans for hiPSC lines with high or low level of XIST mRNA. Only the genes which had ASE data in at least 10 cell lines in both low-XIST and high-XIST female lines are included, excluding the PAR genes. F) A cumulative plot of the percentage of genes in Group 1, Group 2, and Group 3 female hiPSCs based on the fraction of cell lines where the genes escape XCI (ASE > 0.1, FDR < 0.05). All female hiPSC lines with ASE data available are included.

**Figure S3.**
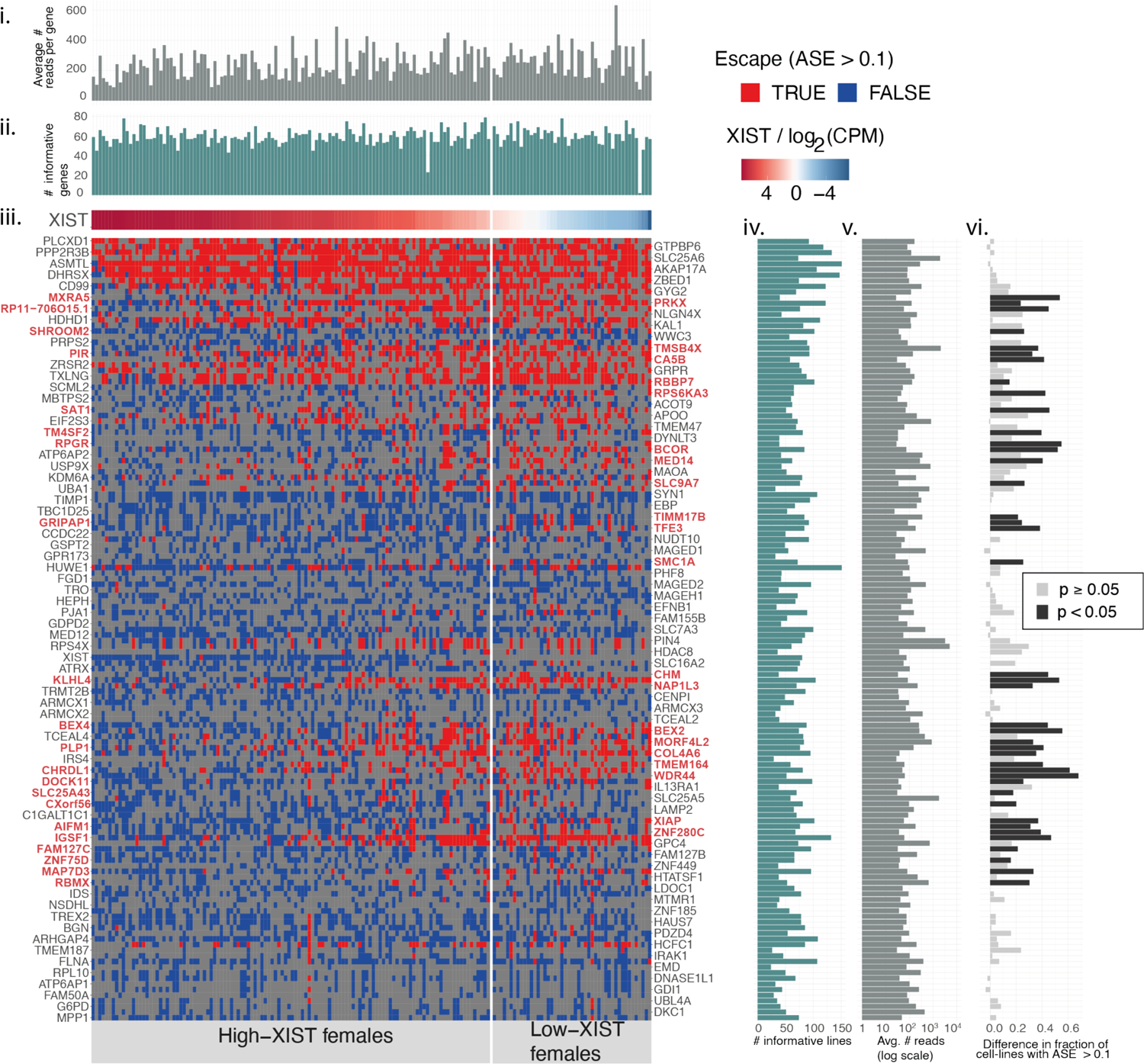
Escape from XCI for 139 X chromosome genes in 165 female lines. The heatmap and bar graphs summarizes the escaping genes (ASE > 0.1, FDR < 0.01) in hiPSCs. The bar graphs above heatmap summarize the average number of sequence reads per gene at an ASE site (i) and number of genes with ASE data available (ii) for each line. The number of informative genes is not significantly associated with the XIST levels (p-value = 0.88). We observed a slight negative linear relationship between the average number of reads per gene at an ASE site and the XIST levels (p-value = 0.02). This is expected given that loss of XIST is associated with higher gene expression from X chromosome. iii) The heatmap displays the 139 genes (rows) that escape (red) or are inactive (blue) for each cell line (columns). Cells in grey color indicate missing information. The low- and high-XIST lines are separated by a vertical line. The genes are ordered by location in X the chromosome and the lines are in decreasing order by XIST level. Names of the genes with significantly (p-value < 0.05, Fisher’s exact test) higher fraction of cell lines with escape in low XIST females than in high XIST females are colored in red. The bar graphs on the right side indicate (iv) the number of lines with ASE data for each gene, and (v) the average number of reads per gene at an ASE site across the lines. vi) A bar graph displaying the difference in the fraction of lines that escape between low and high XIST females. The nominally significant differences (p-value < 0.05) are indicated in black.

**Figure S4.**
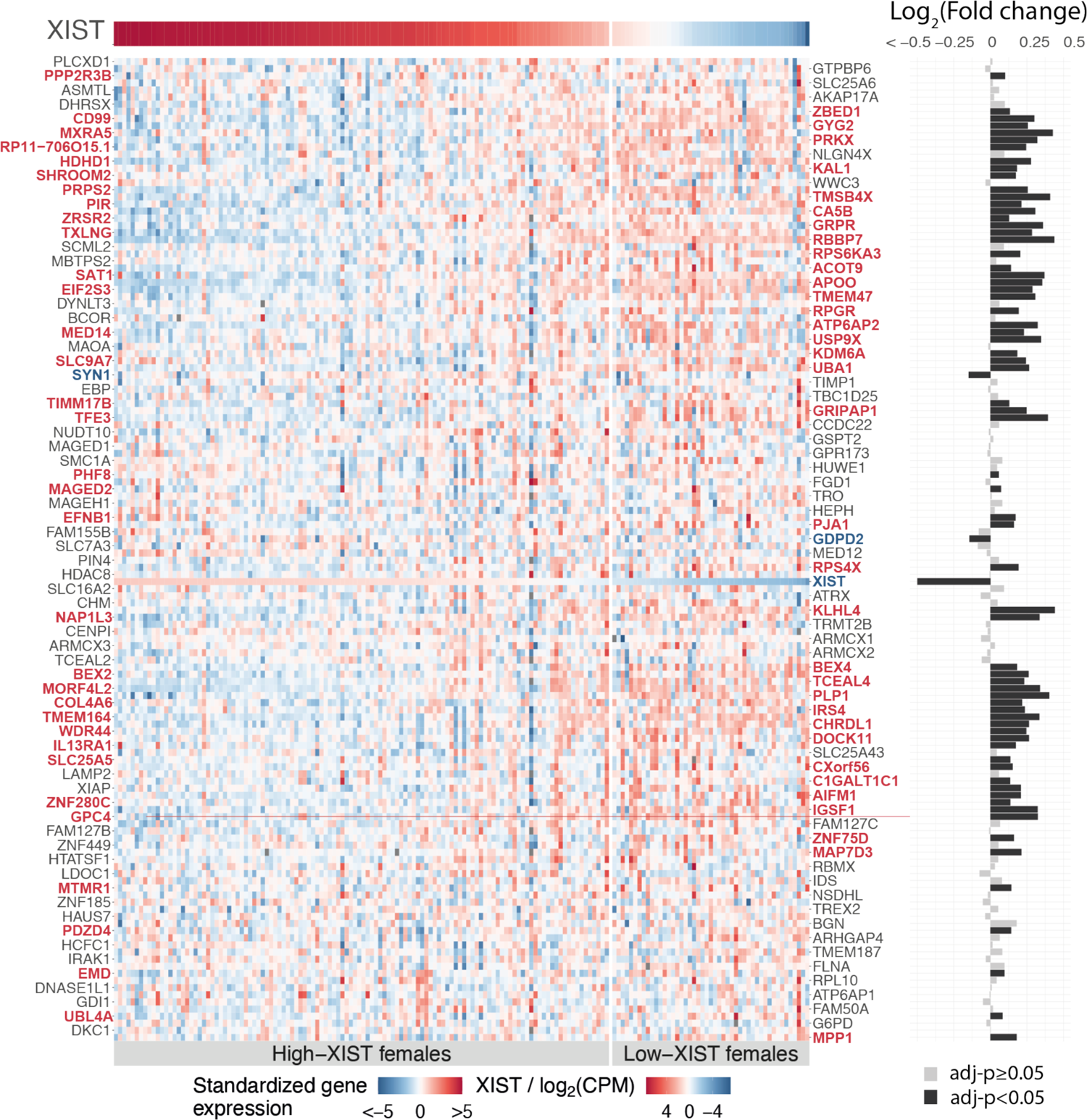
Standardized gene expression for X chromosome genes matched with ENSEMBL identifiers (GencodeV19) with Figure 2B and Figure S4 genes (excluding ENSG00000250349 / TM4SF2 with no data) in 165 female lines (columns). The cell lines are ordered in decreasing order of XIST expression. The low and high XIST lines are separated by a vertical line. The genes (rows) are ordered by location in the X chromosome from top (p-arm) to bottom (q-arm). The bar graph on the right side displays differences in gene expression (log_2_(fold change)) between the low and high XIST lines and the significant differences (adjusted p-value < 0.05) are highlighted in black. Names of the genes with significant difference in their expression between low XIST and high XIST lines are colored in red.

**Figure S5.**
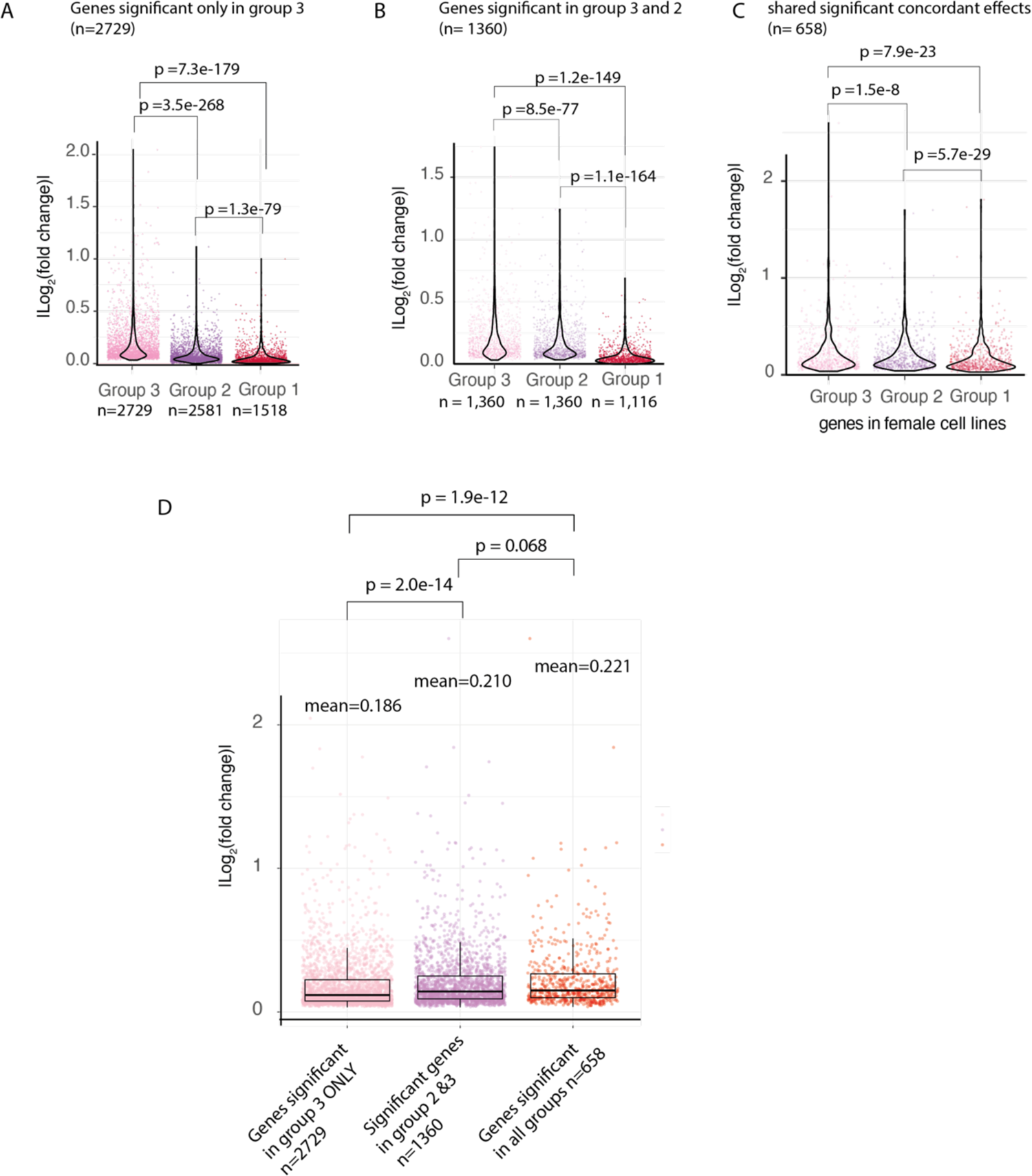
Magnitudes of male-female differences in gene expression are associated dose-dependently to degree of escape from XCI. A-C) Violin plots of absolute log_2_(fold changes) for genes with same direction of effects for male-female difference in autosomal genes in the three groups of female lines. A) Genes that were significant only in group 3 (n=2,729 genes) had concordant effects in 2,581 genes with group 2 and 1,518 genes with group 1. The absolute log_2_(fold changes) were significantly different for the concordant genes (group 3 - group 1 p-value = 7.30e-179, group 3 – group 2: p-value = 3.50e-268, group 2 - group 1: p-value =1.30e-79). Only concordant genes are plotted for each group. B) Genes that were significant in group 2 and group 3 with same direction of effect (n=1,360). The absolute log_2_(fold changes) were significantly different between the groups (group 3 - group 1 p-value = 1.20e-149, group 3 – group 2: p-value = 8.50e-77, group 2 - group 1: p-value = 1.10e-164). Only genes with concordant direction of effect are plotted for group 1 (n=1,116). C) Genes that have significant effects in all female groups with same direction of effect (n=658). The magnitudes of male-female differences differ significantly between the groups (group 3 - group 1: p-value = 7.90e-23, group 3 – group 2: p= 1.50e-08, group 2 - group 1: p-value = 5.70e-29). D) A Tukey style box plot of absolute log_2_(fold changes) of male-female differences in group 3 for significant genes in group 3 only, significant genes in group 2 and group 3 with concordant direction of effect and genes significant in all groups. The means of the absolute log_2_(fold changes) are presented in the plot together with p-values for comparison between the gene groups.

**Figure S6.**
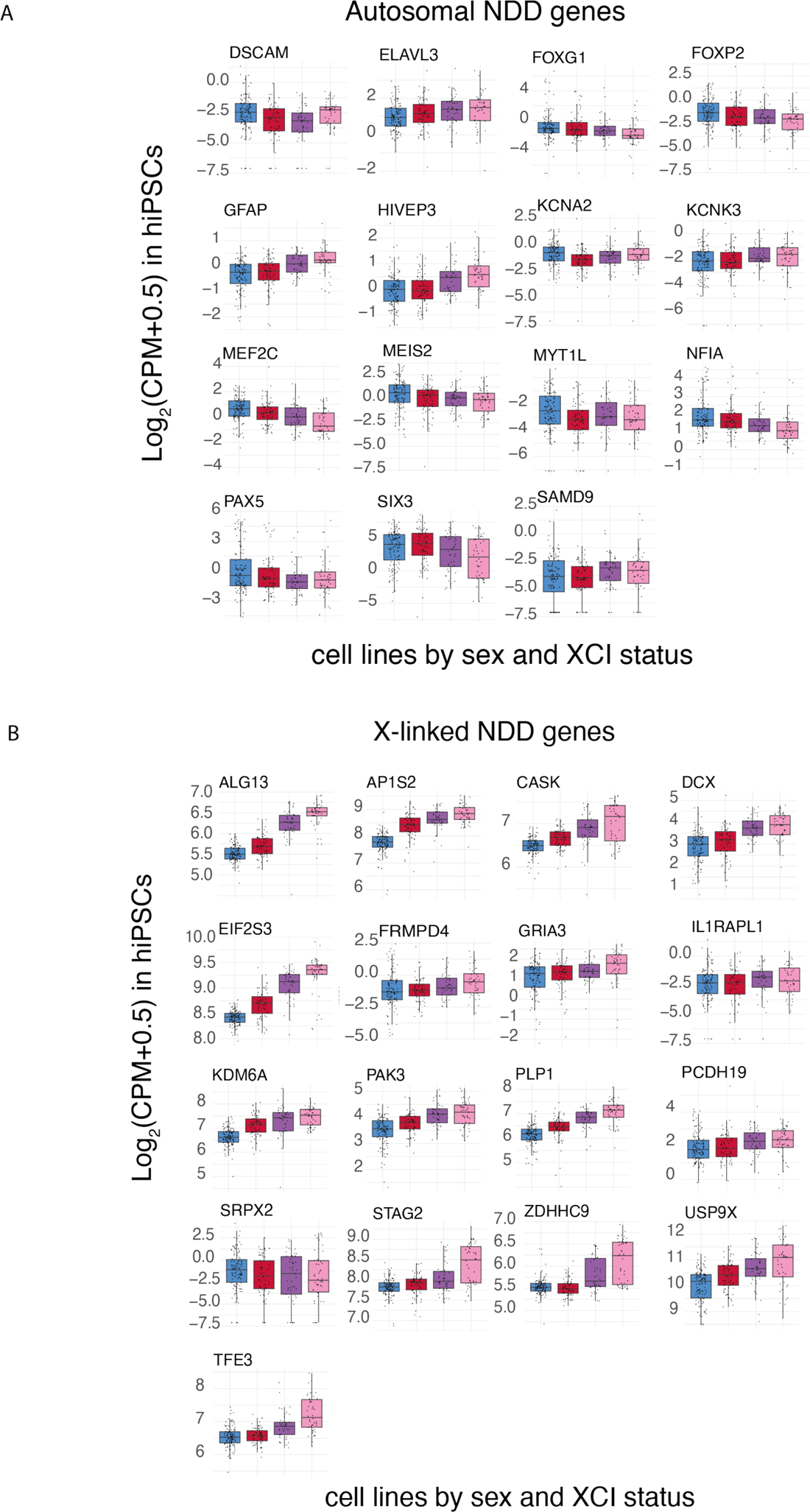
Expression of NDD genes in hiPSCs. A) Expression of autosomal genes in the three groups of female lines and male lines. B) Expression of X chromosome genes in the three groups of female lines and male lines.

**Figure S7.**
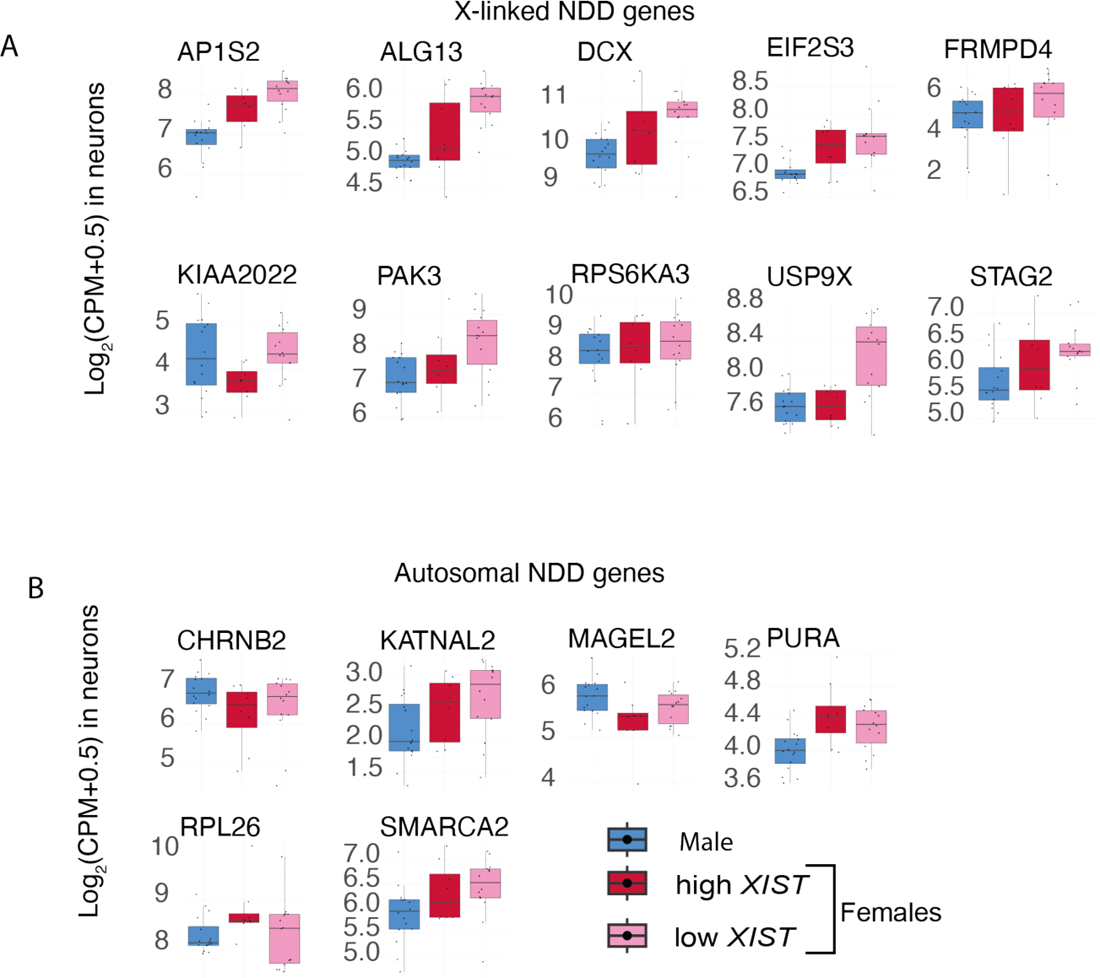
NDD gene expression in hiPSC-derived sensory neurons for A) X-chromosome and B) autosomal genes.

